# Direct voltage imaging of peripheral axons *in vivo* reveals ongoing nociceptor firing in healthy mice

**DOI:** 10.64898/2026.09.19.752889

**Authors:** Efrat Sheinbach, Rachely Butterman, Shaya Lev, Shulamit Baror-Sebban, Alexander M. Binshtok, Yoav Adam

## Abstract

The electrical output of a neuron is generated and conducted along its axon, yet the axon has remained largely inaccessible to direct measurement of membrane potential in intact tissue. Genetically encoded voltage indicators have enabled optical recording of membrane potential dynamics from neuronal cell bodies and dendrites *in vivo*, but direct voltage imaging of identified axons in the living animal has remained largely unexplored. This gap is especially consequential for nociceptive neurons, whose signals are generated and propagated at peripheral terminals far from the soma. Here, we expressed genetically encoded voltage indicators in trigeminal neurons and, using high-speed widefield imaging with holographic targeted illumination, recorded membrane-voltage dynamics directly from corneal nociceptive terminal arbors and their parent axons *in vivo*, in intact tissue. Individual terminals responded to capsaicin and menthol with depolarization accompanied by trains of spiking activity. Unexpectedly, a substantial fraction generated ongoing action-potential-like firing in the absence of an applied stimulus. This firing was sodium-channel-dependent, synchronized across sister terminal branches, and shared its dynamics with the parent axon. Ongoing terminal firing persisted during lidocaine delivery to the trigeminal ganglion, supporting a peripheral origin distal to the ganglion, and was abolished by inhibition of Na_V_1.8 or HCN/Ih conductances. Thus, direct voltage imaging reveals ongoing and spatially coordinated electrical activity within intact peripheral sensory arbors. More broadly, this approach provides access to the generation and integration of neuronal output within fine axonal compartments *in vivo*.

## Introduction

The electrical output of a neuron is generated and propagated along its axon. Yet axons have remained among the least accessible compartments for direct measurement in intact tissue of the living animal. Genetically encoded voltage indicators (GEVIs) report membrane potential optically with millisecond resolution and have enabled *in vivo* recording of action potentials and subthreshold dynamics from neuronal somata and dendrites^1–8^. However, direct voltage imaging of identified fine axons *in vivo* remains limited, leaving the electrical dynamics of these compartments substantially less accessible than those of neuronal somata.

This gap is especially consequential for primary pain-related nociceptor neurons, in which initial detection of noxious stimuli and the generation of a propagating electrical signal occur at the nociceptor terminals ^9–11^. These tiny distal specializations of the peripheral nociceptive axon, embedded in tissue far from their somata in the dorsal root or trigeminal ganglion, generate a graded receptor potential and translate it via voltage-gated channels into action potentials that propagate along the peripheral axons toward the CNS^11,12^. Individual terminals are organized into morphologically complex arbors whose branches differ in length and geometry and converge, through a hierarchy of nodes, onto a parent axon^13–16^. The peripheral arbor therefore constitutes an input–output compartment in which local transduction must be converted into a unitary axonal firing ^17,18^.

Despite the central role of nociceptive endings in detecting noxious signals, most knowledge of nociceptor excitability comes from recordings made far from the terminal: at the soma, along whole nerves, or in dissociated or *ex vivo* preparations ^19–22^. These approaches have been indispensable, but sensory signals originate at the terminal, rather than at the soma or proximal nerve, and physically accessing the terminal risks disrupting the tissue microenvironment that shapes its activity. Optical methods have narrowed the gap: *in vivo* calcium imaging of corneal terminals has resolved where stimulus-evoked signals arise and how they change with inflammation ^17,23^. Calcium signals, however, are an indirect proxy for spiking; they reflect calcium entry through both voltage-gated and transducer channels, as well as complex intraterminal calcium dynamics ^24–26^, and therefore cannot directly report the input-output relation of nociceptive terminals.

We reasoned that the cornea offers a favorable setting to address this: its transparency and accessibility make the fine axons and terminal arbors of trigeminal nociceptive neurons optically accessible in intact tissue^17,23^. We therefore expressed GEVIs in trigeminal neurons and, using a high-speed widefield system combined with holographic targeted illumination, resolved voltage dynamics along axons *in viv*o and evaluated the electrical activity of nociceptive endings.

We first established that individual endings generate voltage responses and action-potential firing during focal application of noxious stimuli. Unexpectedly, we also detected ongoing firing in unstimulated terminals. These events were sodium-channel-dependent, synchronized across sister branches of individual arbors, and shared firing dynamics with the parent axon. Ongoing firing persisted during lidocaine delivery to the trigeminal ganglion, supporting a peripheral origin, and required Na_V_1.8 and HCN/Ih conductances. Thus, direct voltage imaging reveals both ongoing electrical activity and its coordination across the branched architecture of intact peripheral sensory endings, providing access to the transformation of terminal activity into axonal output *in vivo*.

## Results

### Direct *in vivo* voltage imaging resolves electrical activity within corneal nociceptive terminals

To directly measure the electrical activity and output of nociceptive terminals, we expressed the genetically encoded voltage indicator (GEVI) Archon1^27^ in trigeminal ganglion (TG) neurons whose peripheral terminals innervate the cornea^28^. We co-injected Cre-dependent Archon1 (AAV9-hSyn-DIO-Archon1-KGC-EGFP-ER2) with AAV-hSyn-Cre into the TG of adult female C57BL/6 mice (*see Methods,* **Fig. 1a**). Eight to sixteen days later, fluorescently labeled axons and terminal arbors were visible in the cornea (**Fig. 1a**, *inset*). Changes in Archon1 fluorescence were measured through a stabilized eye cup on an upright widefield system equipped with a spatial light modulator for targeted illumination and a fast sCMOS camera ^3,5,29^ (**Fig. 1a**; **Supplementary Fig. 1**). This preparation enabled high-speed (500-1000 frames s^-1^) optical signal detection from small peripheral axons while preserving intact tissue and its innervation. GEVI fluorescence dynamics were analyzed using independent-component analysis (ICA) and expressed in units of the baseline noise standard deviation (σ; *see Methods*).

**Fig. 1.**
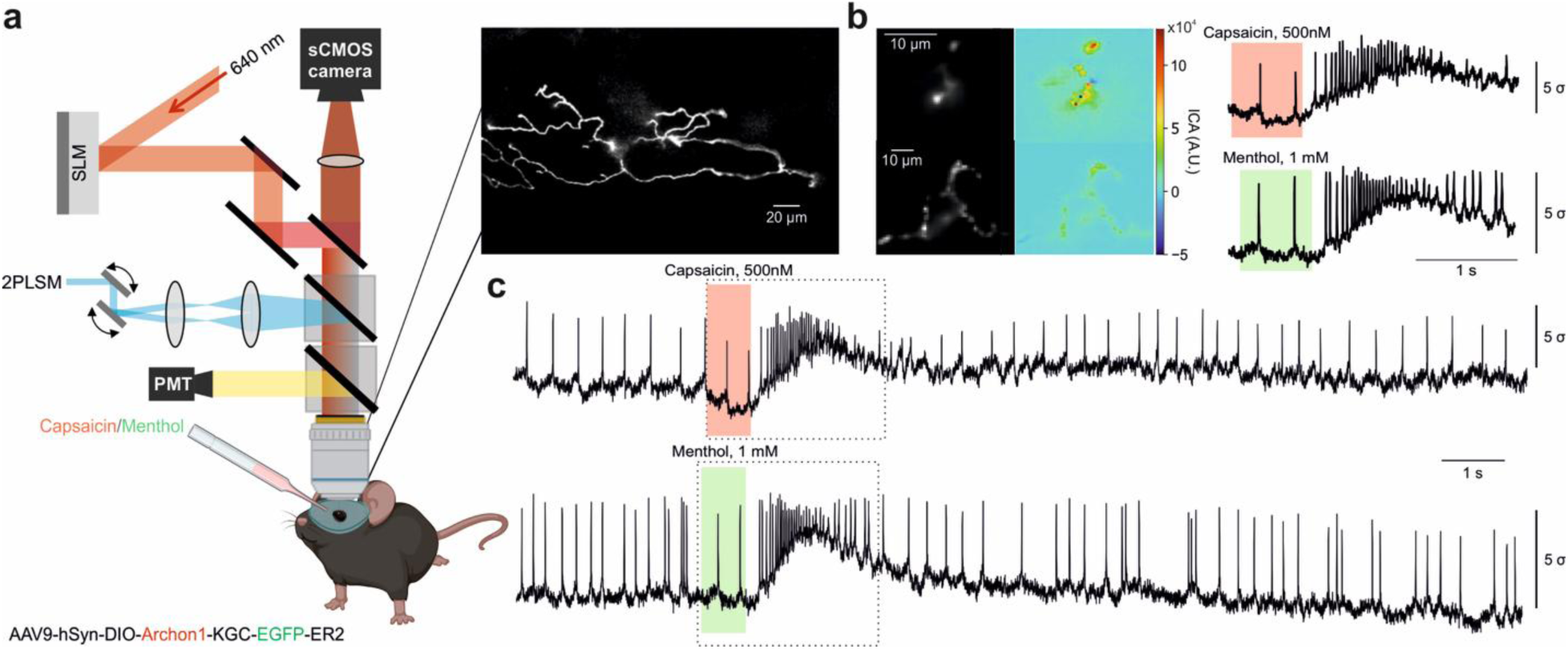
Direct *in vivo* voltage imaging resolves evoked and ongoing electrical activity in corneal nociceptive terminal arbors. (a) Left: schematic of the widefield voltage-imaging system and the Cre-dependent construct AAV9-hSyn-DIO-Archon1-KGC-EGFP-ER2; SLM: spatial light modulation, 2PLSM: two-photon laser scanning microscope. *Right*: representative Archon1/EGFP fluorescence image of a corneal terminal arbor. (b) *Left*: fluorescence images and independent-component (ICA) maps of the representative recorded terminals following the focal application of capsaicin (500 nM, *upper, pink*) or menthol (1 mM, *lower, green*). *Right*: representative voltage traces during focal application of capsaicin (500 nM) or menthol (1 mM). (c) Representative full recordings of the experiments shown in *b,* showing ongoing spike-like events before and after capsaicin (*top*) and menthol (*bottom*) application. Traces are SNR-normalized (*Methods*); amplitudes are in multiples of baseline noise, not absolute voltage. Representative of Capsaicin: n = 5 terminals from N = 5 mice; Menthol: n = 2 terminals from N = 2 mice. The dotted squares indicate the recording times shown in *b*.

We first asked whether the optical signals reported physiological activation of corneal sensory terminals in response to noxious stimuli. Focal application of capsaicin (500 nM) onto individual identified terminal tips via a glass pipette positioned above the corneal epithelium evoked rapid, spike-like optical voltage transients superimposed on a slower, stimulus-associated depolarizing response (**Fig. 1b**). Individual terminals also responded to focal application of menthol (1 mM, **Fig. 1b**). These responses were qualitatively consistent with those we and others reported during current-clamp recordings from trigeminal or dorsal root ganglion neurons exposed to capsaicin or menthol ^25,30,31^.

Unexpectedly, voltage recordings from the capsaicin-sensitive terminals revealed discrete, spike-like voltage transients even before any stimulus was applied (**Fig. 1c**, **Supplementary Fig. 2a–d**). The ongoing activity was observed both in male and female mice. Within terminals, capsaicin-evoked firing occurred at a substantially higher rate than spontaneous firing (**Supplementary Fig. 2e**), whereas the average spike waveform did not differ detectably in half-width or afterhyperpolarization area (**Supplementary Fig. 2a–d, g, h**).

These results establish that GEVI imaging can resolve, in intact tissue, both stimulus-evoked and ongoing voltage events within nociceptive terminal arbors.

### Ongoing voltage events are sodium-channel-dependent and are a common feature of corneal nociceptive terminals

Ongoing activity has previously been described using somatic intracellular recordings in DRGs^32^or extracellular recordings of nerve fibers^33^, and is strongly associated with inflammatory and neuropathic pain states^33–35^. In an *ex vivo* cornea preparation, a proportion of cold-sensitive, but not TRPV1-expressing terminals demonstrate ongoing activity^36^. Thus, the presence of ongoing activity in capsaicin-responsive terminals, as well as in unstimulated terminals in healthy tissue, was unexpected. We therefore next examined whether the ongoing activity we detected reflects locally relevant electrical activity rather than a technical artifact.

Among Archon1-expressing terminals meeting the predefined fluorescence-detection criterion (*see Methods* and **Supplementary Fig. 6**), 110 of 252 (43.7%, 95% CI, 37.6-49.9%, Wilson CI) exhibited ongoing events, which we defined as “active” terminals (*see Methods*). Across the “active” terminals, ongoing events were widespread and heterogeneous in pattern (**Fig. 2a**).

**Fig. 2.**
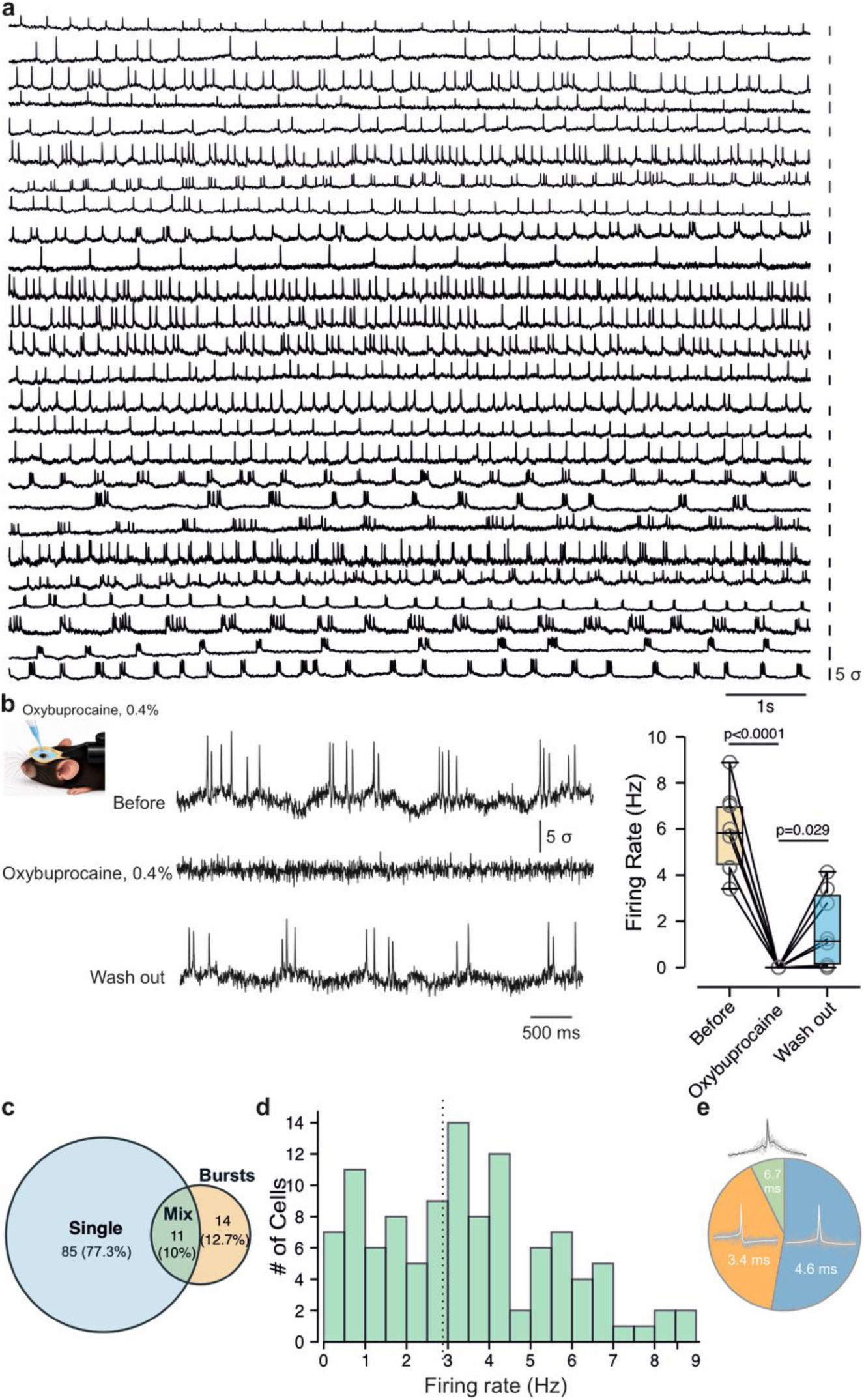
Ongoing voltage events are sodium-channel-dependent action potentials and common in corneal nociceptive terminals. (a) Ongoing-activity traces from many terminals, illustrating heterogeneous firing. (b) *Left*: example traces before, during topical 0.4% oxybuprocaine, and after wash-out. *Right*: Box plots with individual values of firing rate in terminals before, during application of oxybuprocaine, and after wash-out; lines connect the same terminal. n = 8 terminals from N = 8 mice; a repeated-measures (RM) one-way ANOVA with post-hoc Bonferroni. (c) Venn diagram depicting firing-pattern classes among active terminals: Single 77.3% (85/110), Burst 12.7% (14/110) and Mix 10% (11/110); n = 110 terminals; N = 42 mice. (d) Distribution of ongoing firing rates across active terminals (n = 110, N = 42 mice); dashed line outlines the mean of 3.49 ± 2.15 Hz. (e) Mean spike waveforms grouped by k-means cluster; labels indicate mean half-widths (*see Supplementary Fig. 6*).

To test whether these ongoing voltage deflections represent action potentials (APs) mediated by voltage-gated sodium channels (Na_V_), we applied the broad-spectrum voltage-gated sodium-channel blocker oxybuprocaine (0.4% ^37,38^) topically to the eye. We previously demonstrated that *in vivo* application of 0.4% oxybuprocaine to the mouse cornea inhibited capsaicin-induced calcium signals in the terminal fibers ^17^. Oxybuprocaine abolished ongoing firing, which recovered after washout (**Fig. 2b**), identifying these events as APs.

The ongoing AP firing fell into three temporal classes defined by an inter-spike-interval mixture model: single-spike (∼77%, 85 of 110), mixed (∼10%, 11 of 110), and burst (∼13%, 14 of 110 active terminals) (**Fig. 2c**; Methods). Active terminals fired at rates broadly distributed around a mean of 3.49 ± 2.15 Hz, ranging from 0.04 to 8.9 Hz, with a median of 3.37 Hz (**Fig. 2d**).

The average AP waveforms of active terminals were heterogeneous: unsupervised clustering of per-terminal waveforms identified three AP clusters differing in half-width (cluster 1, n = 58, half-width 4.58 ms; cluster 2, n = 44, 3.38 ms; cluster 3, n = 8, 6.72 ms; **Fig. 2e**; **Supplementary Fig. 3**). The clusters differed in firing rate and afterhyperpolarization (**Supplementary Fig. 3c-e**), indicating heterogeneity in the waveform and temporal organization of ongoing activity.

Next, we examined whether the ongoing activity was pathological, specifically whether viral expression of Archon1 had impaired terminals and driven ectopic firing rather than revealing a native property of healthy terminals. Corneal instillation of capsaicin evoked robust forelimb eye-wiping in mice expressing AAV9-Archon1-EGFP (before vs capsaicin p = 0.0012), a response indistinguishable from that of mice expressing AAV9-EGFP alone (between-group p > 0.9; **Supplementary Fig. 4**). Archon1 expression, therefore, did not detectably alter capsaicin-evoked nocifensive behavior, arguing against GEVI-induced dysfunction as the source of ongoing firing.

Moreover, the ongoing activity was not driven by local heating of the terminals from the excitation laser (**Supplementary Fig. 5**). Ongoing firing was also reproduced using a spectrally and mechanistically distinct indicator, Ace-mNeon2^4^, and was uncorrelated with GEVI expression, as measured by terminal brightness, further suggesting that it was not an artifact of the indicator overexpression or of signal intensity (**Supplementary Fig. 6a-d**).

Together, converging pharmacological, behavioral, optical, and cross-indicator controls indicate that ongoing voltage events reflect sodium-channel-dependent AP firing in a substantial fraction of corneal nociceptive terminals in otherwise-normal mice.

### Ongoing firing is coordinated across sister terminal trees and their parent axon

Corneal nociceptive neurons develop morphologically complex terminal arbors, in which multiple axonal terminal branches of varying length and geometry converge through successive nodes onto a parent axon^13,14,28,39^. This architecture raises the question of how ongoing firing is integrated across the branches of a single tree.

In a subset of recordings, two or more adjacent terminal “sister” branches of the same terminal arbor fell within a single field of view and were imaged simultaneously (**Fig. 3**). Although the sister terminals were spatially distinct in the fluorescence image, independent-component analysis (ICA) assigned their activity to a single temporal component: the branches carried near-identical, time-locked spike trains and thus behaved as a single spiking unit (**Fig. 3a**). To quantify this synchrony, we extracted independent ROIs from each sister branch and compared their spike trains (**Fig. 3b**). Confirming the single-component ICA result, the cross-correlogram peaked at zero lag with a full width at half maximum (FWHM) of about 2 ms, which is as narrow as the acquisition resolution allowed, and showed no consistent lead or lag (**Fig. 3c**). Thus, spatially distinct sister branches behave electrically as a coordinated spiking unit at the temporal resolution of our recordings.

**Fig. 3.**
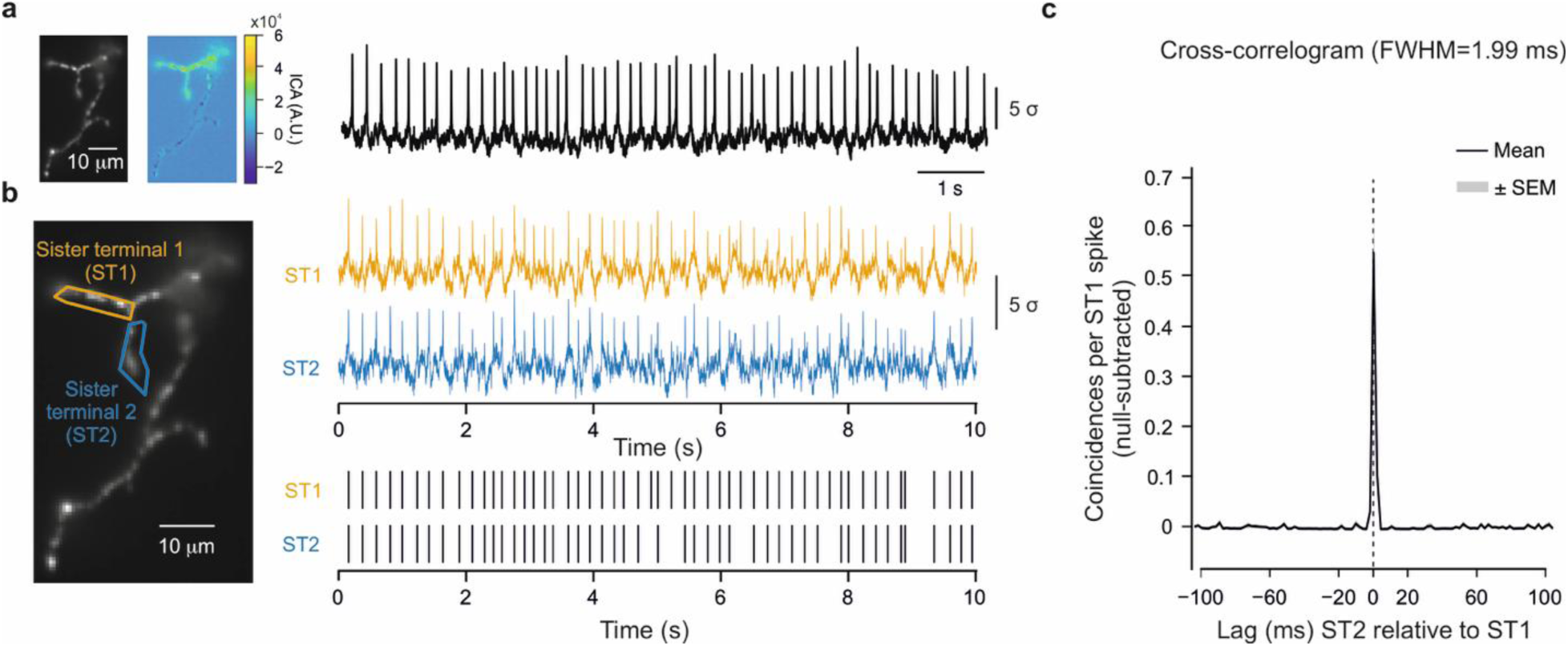
Sister terminals of a terminal arbor fire in tight synchrony **(a)** *Left*: fluorescence image of the arbor and the corresponding independent-component (ICA) map. *Right*: the extracted voltage trace of the common independent component, showing ongoing spike-like events. The two sister branches are not separated into distinct components, indicating that their activity is essentially inseparable by ICA. **(b)** *Left*: fluorescence image with two manually defined regions of interest (ROIs), sister terminal 1 (ST1, *orange*) and sister terminal 2 (ST2, *blue*). *Right*, top: simultaneously recorded voltage traces from ST1 (*orange*) and ST2 (*blue*); bottom: corresponding spike rasters for ST1 and ST2, showing near-coincident firing. **(c)** Pairwise cross-correlogram (mean ± SEM across pairs) of null-subtracted coincidences per ST1 spike as a function of lag (ST2 relative to ST1). The correlogram peaks sharply at zero lag (full width at half-maximum 1.99 ms) with no consistent lead or lag, indicating near-simultaneous firing with no detectable directional offset. Data are from n = 5 sister-terminal pairs from N = 5 mice.

However, the spatial resolution and signal-to-noise of Archon1 limited our imaging to only a few terminal branches. To determine how ongoing firing is integrated within the arbors, along the terminal tree and the parent axon, we needed to record from multiple terminal structures and compare these recordings with those from the parent axon. Therefore, for these experiments, we used a cell-filling version of the GEVI Ace-mNeon2 ^4^, by fusing it with the N-terminal Lucy-Rho tag^40^, previously shown to improve the dendrite expression of the GEVI Voltron2^41^ (AAV-CAG-DIO-LR-Ace-mNeon2-TS-ER2; **Fig. 4a**, *Methods*). The brighter fluorescence and stronger expression of LR-AcemNeon2 in fine, deep processes improved spatial resolution, allowing us to consistently identify a parent axon and associated terminal arbors with multiple “sister” terminal fibers within a single field (**Fig. 4a,b, Supplementary Movie 1**). Because the terminal arbors and the parent axon were at different tissue depths, they were recorded separately. That is, activity within a field of view was simultaneous, but activity across fields was not. We therefore could not assess synchrony between the two arbors or between terminals and the axon; instead, we compared their firing dynamics.

**Fig. 4.**
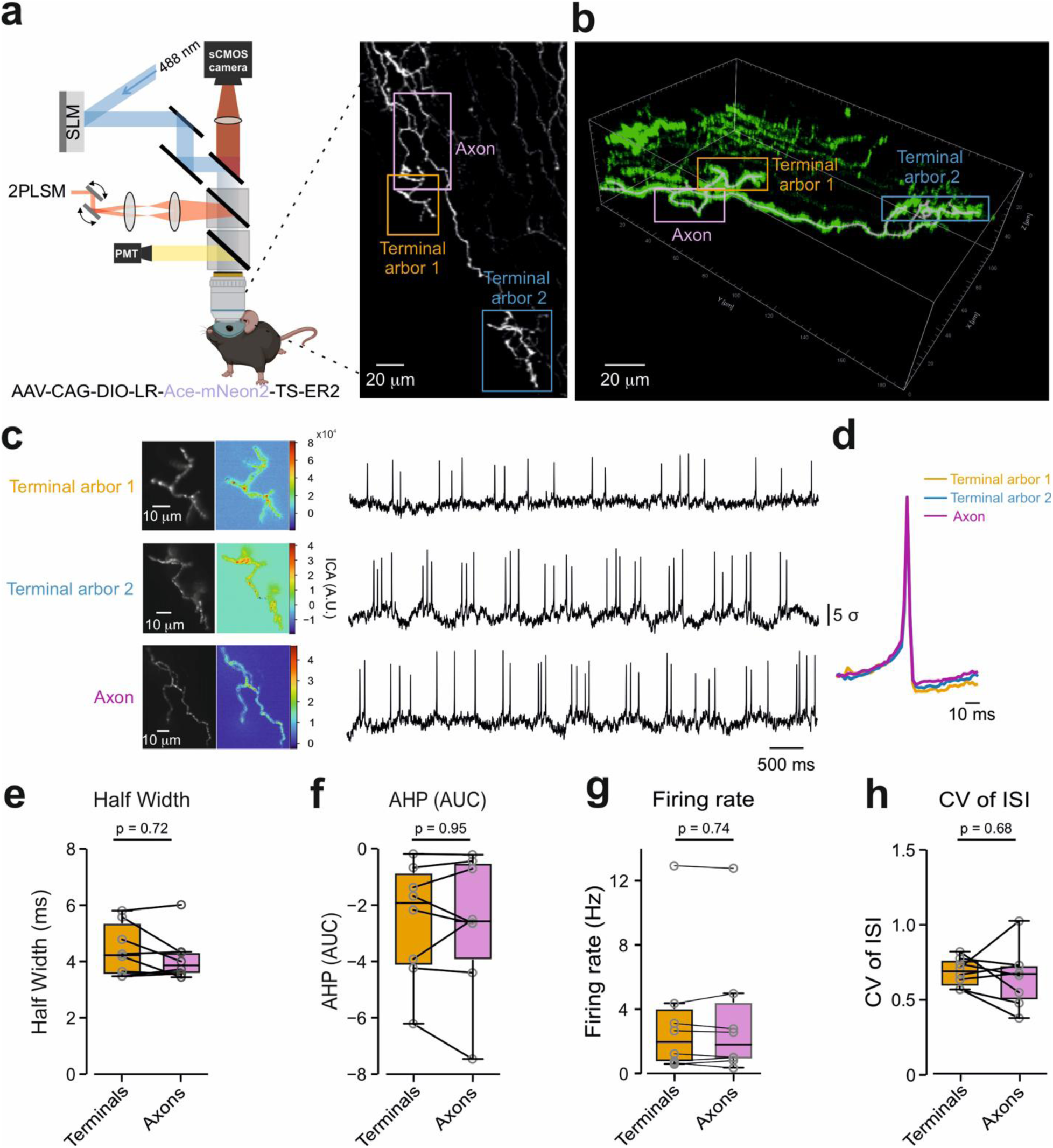
Ongoing firing is coordinated across sister terminal trees and shares its dynamics with their parent axon. (a) *Left*: schematic of the recording setup for Ace-mNeon2 detection and Cre-dependent construct AAV-CAG-DIO-LR-Ace-mNeon2-TS-ER2. *Right*: fluorescence image identifying a parent axon and two terminal arbors (arbor 1, *orange*; arbor 2, *blue*). (b) Three-dimensional reconstruction of the same terminal tree (*see also Supplementary Movie 1*). (c) Voltage traces from terminal arbor 1, terminal arbor 2, and the parent axon. (d) Overlaid mean spike waveforms from the two arbors and the axon. (**e–h**) Paired comparison of terminal arbors vs their parent axons: (**e**) half-width; (**f**) afterhyperpolarization (AHP) area; (**g**) firing rate; (**h**) coefficient of variation of interspike interval (CV of ISI). Box plots with individual pairs; Wilcoxon signed-rank; n = 8 terminal–axon pairs from N=8 mice.

Consistent with the sister-terminal synchrony observed with Archon1 (**Fig. 3**), independent-component analysis of the Ace-mNeon2 recordings resolved the activity of adjacent sister terminal branches of the same arbor into a single temporal component, indicating that the branches generated near-identical, time-locked spike trains and operated as a single spiking unit (**Fig. 4c**). When we compared firing at separate ROIs from each sister branch (**Supplementary Fig. 7a, b**), the cross-correlogram peaked sharply at zero lag (FWHM of 2.49 ms) with no consistent lead or lag (**Supplementary Fig. 7b**).

Comparing terminal arbors with their parent axons across recordings showed that the average spike waveforms recorded at the two arbors and at the axon were closely superimposable (exemplified in **Fig. 4d**), with no systematic differences in spike half-width and afterhyperpolarization area (**Fig. 4e, f**). Importantly, firing rate and interspike interval (ISI) variability were also similar across the paired terminal and parent-axon recordings (**Fig. 4g, h**).

### Ongoing terminal firing persists during lidocaine delivery to the trigeminal ganglion

The observations that terminal and axonal rates matched and that waveforms were nearly identical across sites argue against independent summation of branch-generated APs, because in that case the axon’s event rate would be expected to exceed that of any single branch. Thus, we hypothesized that the ongoing activity is generated locally and synchronized across branches, or that it is generated in a common region, such as somata within the TG, and propagates antidromically to the recorded sites.

To test whether ongoing terminal activity requires activity originating at or proximal to the TG soma, we implanted a cannula into the TG using the coordinates of the original viral injection and delivered 2% lidocaine to the somata of primary sensory neurons, as it was previously demonstrated that intra-TG injection of 2% lidocaine substantially suppressed trigeminal afferent signaling *in vivo* in rats^42^. The lidocaine solution was mixed with Evans Blue, which served as a delivery marker. We then recorded ongoing activity at the terminals before and 3 minutes after lidocaine delivery to the TG (**Fig. 5a**; *Methods*). After imaging, the TG was extracted and inspected for blue coloration to confirm that the injectate reached the ganglion (**Fig. 5a**). In agreement with the hypothesis of local generation of ongoing activity, ongoing terminal firing persisted after lidocaine delivery to the TG (**Fig. 5b**). Firing rate, CV of ISI, spike half-width, afterhyperpolarization area, and the inter-spike-interval distribution (**Fig. 5c-e**) were not detectably changed, and the average waveform was unchanged (**Fig. 5f-h**) ^42^. These observations argue against the TG soma or more proximal sites as the source of ongoing terminal activity and support an origin in the peripheral branch distal to the ganglion.

**Fig. 5.**
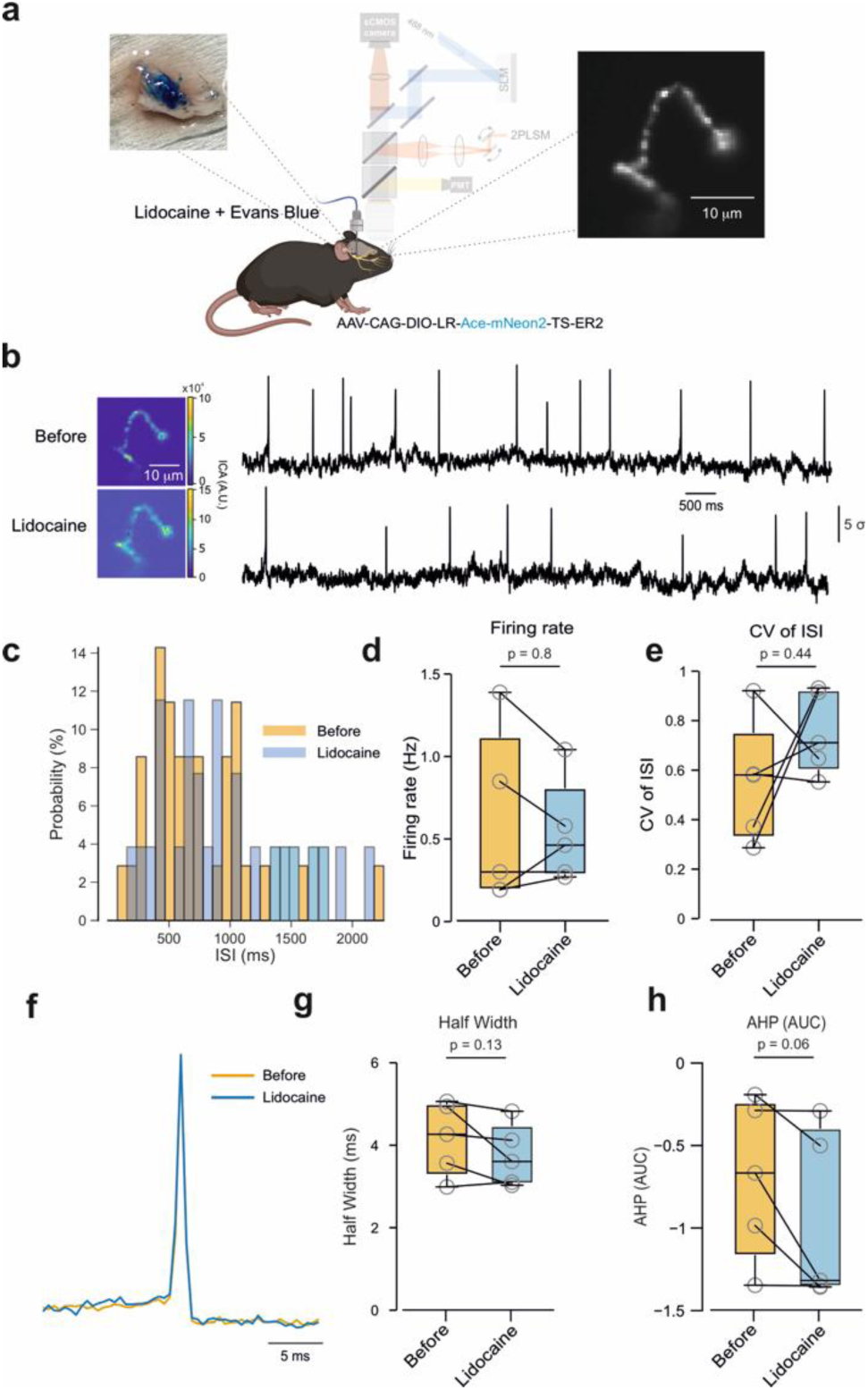
Ongoing terminal firing persists during lidocaine delivery to the trigeminal ganglion. (**a**) Schematic of TG cannulation and delivery of 2% lidocaine with Evans Blue (delivery marker); insets, *left*: delivery validated by post-hoc blue coloration of the extracted TG by Evans Blue; *right*: a representative terminal arbor. (**b-c**) Example traces before and after TG lidocaine delivery, with fluorescence/ICA maps (**b**) and histograms of the probability distribution of interspike intervals (**c**) before (*orange*) and after TG lidocaine delivery (*blue*). (**d–e**) Paired comparisons of (**d**) firing rate and (e) CV of ISI before and after the application of 2% lidocaine to the TG. Wilcoxon signed-rank test; n = 5 terminals from N = 5 mice. (f) Overlaid mean waveforms of AP from the example shown in *b*. (**g-h**) Paired comparisons of half-width and AHP area. Box plots with individual terminals; Wilcoxon signed-rank test; n = 5 terminals from N = 5 mice.

### Ongoing terminal firing requires Na_V_1.8 and HCN/Ih conductances

Finally, we aimed to identify the conductances underlying the ongoing activity in the peripheral branches. Ongoing activity is expected to require inward current for electrogenesis, a mechanism for recovery between spikes, and a membrane dynamic that permits sustained excitability. Sodium Na_V_1.8 carries a slowly inactivating, tetrodotoxin-resistant sodium current that supports repetitive AP firing in nociceptors and underlies the majority of the current underlying AP in nociceptive axons^43,44^. Indeed, topical application of the selective Na_V_1.8 inhibitor suzetrigine^45,46^ abolished ongoing firing in all recorded terminals (**Fig. 6a**), indicating that Na_V_1.8 is required for ongoing activity.

**Fig. 6.**
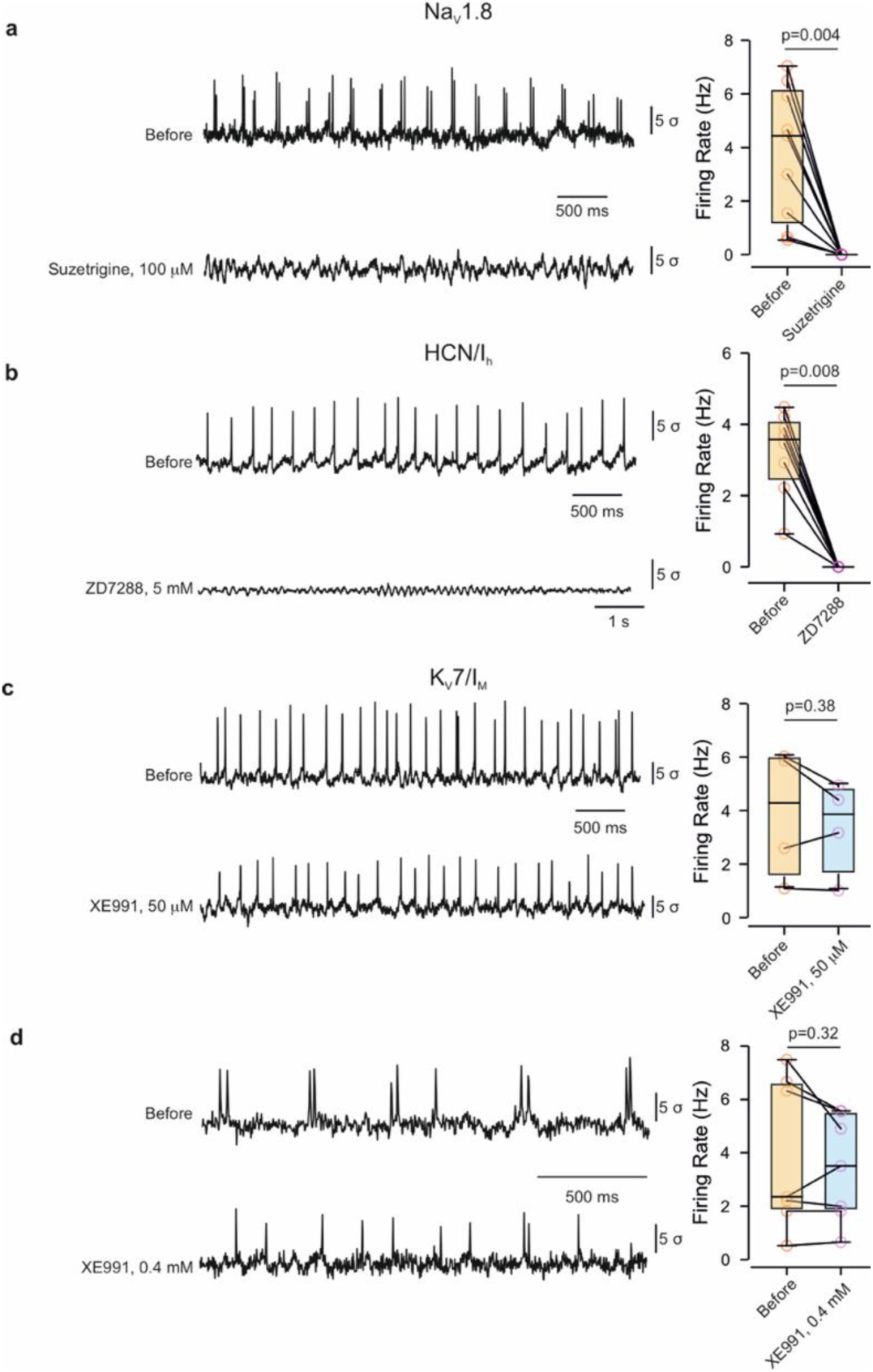
Ongoing terminal firing requires Na_V_1.8 and HCN/Ih currents but not Kv7. **a.** *Left*: representative ongoing-activity traces from the same terminal before (*top*) and during topical application of the selective Na_V_1.8 inhibitor suzetrigine (100 µM, *bottom*); *Right*: firing rate before versus during suzetrigine (box plots with individual terminals; lines connect the same terminal). Suzetrigine abolished ongoing firing in all recorded terminals. n = 9 terminals from N = 9 mice, Wilcoxon signed-rank test. **b.** Same as *a* but for HCN/Ih channel blocker, ZD7288 (5 mM). ZD7288 abolished ongoing firing in all recorded terminals. n = 8 terminals from N = 8 mice; Wilcoxon signed-rank test. **c-d.** Same as *a* but for K_V_7 channel blocker, XE991: 50 µM (**c**) and 0.4 mM (**d**). XE991 produced no detectable change in ongoing firing. For **c,** n = 4 terminals from N = 4 mice; Wilcoxon signed-rank test. For **d,** n = 7 terminals from N = 7 mice; Wilcoxon signed-rank test.

Na_V_1.8, however, is available at relatively depolarized potentials. Therefore, we next examined currents active in the subthreshold range whose activation or inhibition could depolarize the membrane sufficiently to activate Na_V_1.8. The hyperpolarization-activated current Ih (HCN channels) is active at rest, depolarizes the membrane, mediates the post-hyperpolarization sag and rebound that follow each spike, and underlies pacemaker-like rhythmic firing in many excitable cells^47^. We therefore hypothesized that its activation may also be required for the generation of ongoing activity. Application of the HCN/Ih blocker ZD7288^48^ eliminated ongoing firing in all recorded terminals (**Fig. 6b**), consistent with a requirement for Ih in sustaining repetitive terminal activity.

Another potential candidate, acting in the subthreshold range and expressed in nociceptive terminals, is the K_V_7/M current^49^. K_V_7/M channels carry a slowly activating, non-inactivating potassium current that is active near resting potential and acts as a brake on repetitive firing ^50,51^; their inhibition depolarizes the membrane and increases excitability, and loss of K_V_7 function is sufficient to generate ectopic and spontaneous firing in nociceptors^52^. We therefore expected that inhibition of K_V_7 would increase ongoing activity. Contrary to this expectation, the K_V_7 blocker XE991^17,53^ produced no detectable change in ongoing firing when applied at 50 μM (**Fig. 6c**) or even at a higher concentration of 0.4 mM (**Fig. 6d**). We previously demonstrated that 50 μM XE991 applied *in vivo* to the cornea was sufficient to increase capsaicin-induced sodium-dependent responses in nociceptive terminals^17^.

These results identify NaV1.8 and HCN/Ih conductances as necessary for ongoing terminal firing and suggest a model in which NaV1.8 supports regenerative spike generation while Ih contributes to subthreshold excitability and repetitive firing.

## Discussion

Direct access to membrane voltage in fine axons and their terminal arbors has remained a major limitation for studying neuronal output in intact tissue. Here, genetically encoded voltage indicators allowed us to resolve action-potential dynamics directly from peripheral sensory terminal arbors and their parent axons *in vivo*. This revealed ongoing sodium-channel-dependent firing in unstimulated terminals, coordination of firing across sister branches, and similar firing dynamics in terminal arbors and their parent axons. Ongoing activity persisted during lidocaine delivery to the trigeminal ganglion and was abolished by inhibition of Na_V_1.8 or HCN/Ih conductances. Together, these observations reveal an electrically active and highly coordinated peripheral axonal compartment whose dynamics can now be measured directly *in vivo*.

Ongoing firing is closely linked to pathology; however, nociceptive terminals appeared healthy following expression of various GEVIs, and viral expression of Archon1 did not produce hyper- or hyposensitivity in response to a natural noxious stimulus (**Supplementary Fig. 4**). Ongoing firing events were not produced by heating or imaging itself, because firing rate was invariant across a threefold range of excitation-laser power, ruling out an illumination- or heating-driven origin (**Supplementary Fig. 5**). They were not an indicator-specific optical artifact: the phenomenon reappeared with a spectrally and mechanistically distinct GEVI, Ace-mNeon2 (**Fig. 4**). The two indicators reported different proportions of active terminals: 43.7% with Archon1 and 28.9% with Ace-mNeon2, but this is most simply explained by differences in sensitivity, expression depth, and sampling rather than by biology. The firing rate was uncorrelated with terminal brightness for both indicators, so active and inactive terminals could not be distinguished by signal intensity (**Supplementary Fig. 6**). Ongoing firing was abolished by a sodium-channel blocker and recovered on washout (**Fig. 2b**), the defining signature of action potentials.

Human microneurography recordings show ectopic activity in neuropathic and inflammatory states but not in healthy nerves^33,35^. The reason for this discrepancy could be that microneurography accesses the axon several centimeters proximal to the endings, where terminal-generated activity may be filtered at branch points before it reaches the electrode. Another important difference is the recorded tissue. Corneal afferents comprise functionally distinct populations, including polymodal (TRPV1-expressing) nociceptors and cold-sensitive (TRPM8-expressing) thermoreceptors. The latter, but not TRPV1-expressing fibers, can exhibit ongoing activity under physiological conditions^36^. Importantly, we demonstrated that ongoing activity is also present in capsaicin-sensitive terminals (**Fig. 1**, **Supplementary Fig. S2**).

If a large proportion of nociceptive terminals fire continuously, why is the healthy eye not in constant pain? It could be that ongoing peripheral activity and pain perception are separated by one or more filters. Use-dependent conduction failure at the sensory-neuron T-junction acts as a low-pass filter for C-type afferents^54,55^, and events may also fail at convergence points before reaching the parent axon^56^. However, we show that terminal and parent-axon firing rates matched (**Fig. 4**), which, if general, implies that ongoing events are not strongly filtered before the parent axon and shifts the locus of filtering more centrally, to the T-junction or the CNS. In the CNS, the rate of ongoing activity may fall below that needed to recruit spinal or brainstem nociceptive circuits; the active population may be sparse and asynchronous at the network level; and downstream adaptation, inhibitory gating, and descending modulation could each discard a tonic input. More generally, the nociceptive code depends more on changes in baseline, synchrony, pattern, and population recruitment than on the mere presence of spikes^57^.

Ongoing peripheral activity may also have local functions independent of conscious sensation. Action potentials can promote Ca²⁺ entry and tonic neuropeptide release from sensory endings, raising the possibility that basal firing contributes to tonic neuro-effector signaling. This possibility may be particularly relevant in the cornea, where sensory innervation contributes to epithelial homeostasis and ocular-surface regulation^58,59^.

A striking feature of these recordings is that the terminal tree has many branches, capable of firing, yet the axon’s ongoing rate matches that of a single terminal rather than the sum of its branches (**Fig. 4**). How might activity from many terminals be integrated without summing? One possibility is that the branches do not fire independently at all: if sister terminals are synchronized (**Fig. 3a**), their near-coincident spikes could interact or collide at the convergence nodes, so that a synchronous volley across the tree is read out as a single propagating event on the parent axon. A second, non-exclusive possibility is imposed by the geometry of the arbor itself. Our modeling of the corneal terminal tree predicted that, for brief action-potential-like events, activating one branch or all branches yields the same output at the parent fiber, because refractoriness and sodium-channel inactivation at each convergence point allow only one event to pass and transiently render the converging branches unexcitable; summation emerges only for slow, sustained depolarizations, not for discrete spikes ^39^. Ongoing spontaneous firing, composed of such discrete events, would therefore be expected to integrate sub-additively, with the terminal tree behaving as a coincidence-gated single unit rather than a summing junction. This framework is consistent with the coordinated branch activity and terminal–axon firing dynamics observed here. An important consequence of this organization is that ongoing and stimulus-evoked spikes must share the same convergence points and parent axon. Whether ongoing activity alters the gain or propagation fidelity of subsequently evoked signals is therefore an important question for future experiments.

What produces repetitive firing within a terminal? Our pharmacological experiments demonstrate that blocking either Na_V_1.8 or HCN/Ih channels prevents ongoing activity. One parsimonious model is that Na_V_1.8 provides regenerative inward current once threshold is reached ^43,60^, whereas HCN/Ih contributes to the subthreshold membrane dynamics that permit repetitive firing^47,61^. The organelle-rich, high-resistance terminal branch, with its short space constant^39^, would tend to compartmentalize and amplify small local currents, lowering the charge needed to reach threshold. In this picture, the absence of a detectable XE991 effect indicates that K_V_7-mediated restraint is not required to permit ongoing firing under the conditions tested.

Na_V_1.9 is another candidate contributor because its persistent TTX-resistant current operates near resting potential and can amplify subthreshold depolarization^62^. Resolving its contribution will require selective acute manipulations that were not available in the present study Several limitations constrain our conclusions. The pharmacological agents we used target only the channels we inferred from known action potential physiology, so other factors may contribute to spontaneous activity. Moreover, the method has finite spatial and temporal resolution and a signal-to-noise-limited detection threshold referenced to each recording’s noise, so slow, small, or low-rate events and the smallest, dimmest terminals are undersampled. Finally, all recordings were made under anesthesia, which alters membrane excitability, so the rates reported here need not match those of the awake animal. The findings are so far confined to the cornea; their generalization to nociceptive terminals in the skin, joints, or viscera remains to be tested.

Within these bounds, the electrical output of nociceptive nerve endings and axons can now be observed directly in the intact animal. Direct voltage imaging therefore reveals an electrically active and highly coordinated peripheral axonal compartment *in vivo*. Ongoing firing across terminal branches, its representation in the parent axon, and its dependence on Na_V_1.8- and HCN/Ih-sensitive conductances establish a framework for examining how electrical signals are generated and integrated within intact sensory arbors. A branched peripheral sensory ending is not merely a collection of independent transducers; it behaves as a coordinated electrical compartment, and we can now watch that compartment operate *in vivo*. More broadly, extending voltage imaging into fine axons and their terminal structures provides direct experimental access to neuronal output at a compartment that has remained largely inaccessible in the living animal

## Methods

### Animals

Adult (8–10 weeks, 16–20 g) female and male C57BL/6JOlaHsd mice were used. Animals were housed in specific-pathogen-free conditions under a 12-h light/dark cycle at 23 ± 2 °C with controlled humidity, environmental enrichment, and ad libitum food and water, and were randomly allocated to experimental groups. All procedures were approved by the Ethics Committee of the Hebrew University of Jerusalem (protocol MD-21-17909) and conducted in accordance with institutional and national guidelines.

The ongoing activity was observed in both female and male mice. The quantification and pharmacological experiments were performed only on female mice.

### Viral constructs and GEVI expression

AAV serotype 9 vectors were used: AAV-hSyn-DIO-Archon1-KGC-GFP-ER2 (Addgene #115893)^27^, AAV-CAG-DIO-LR-Ace-mNeon2-TS-ER2, and AAV-CAG-DIO-LR-pAce-mNeon-TS-ER2. Because the GEVI cassettes are Cre-dependent (DIO/FLEX), a Cre-expressing virus (AAV-hSyn-Cre) was co-injected; Cre and GEVI viruses were mixed 1:4 (Cre: GEVI) and aliquoted as 4-µl working stocks.

### Stereotaxic viral injection into the trigeminal ganglion

The expression of GEVI at the corneal terminals was achieved by a viral infection of the trigeminal ganglion (TG) cell bodies via stereotaxic injection to the V1 area, as we previously detailed^63^. In short: seven- to nine-week-old mice were anesthetized either with isoflurane (2% induction, ∼0.4% maintenance via nose cone) or with an intraperitoneal ketamine–medetomidine (Domitor)-saline mixture (1:1:8, 9 µl/g). Meloxicam (5 mg/kg s.c. in 0.3 ml saline) was given preoperatively, and ophthalmic ointment (Duratears) was applied to both eyes. After depilation and a ∼1.5-cm midline incision, a hole was drilled at coordinates targeting the TG (AP +0.4 mm; ML ±1.35 mm; DV −6.4 mm). Virus was delivered with an UMP3 microinjection pump (World Precision Instruments) and a 10-µl syringe: 1700 nl per TG for Archon1, 500 nl for Ace-mNeon2, and 1000 nl for pAce-mNeon. The needle was left in place for 5 min before slow retraction. A custom titanium head-plate was affixed with dental cement, and the incision closed with cement (C&B Superbond, Sun Medical). Imaging was performed 8–16 days after injection.

### In vivo imaging setup and acquisition

Imaging used an upright Bergamo microscope body (Thorlabs) equipped with Olympus XLUMPlanFL N 20×/1.00 NA water-immersion objective, widefield epifluorescence module and a custom holography module (Thorlabs) incorporating a spatial light modulator (EXULUS-HD2, Thorlabs, **Supplementary Fig. 1**). Excitation lasers were a 120-mW CW 488-nm laser (Coherent) for Ace-mNeon, and a 1-W CW 639-nm laser (CNI) for Archon1. Both were fiber-coupled, expanded, collimated, patterned by the SLM, and relayed to the objective back aperture; the zeroth order was blocked in an intermediate image plane. Maximum powers at the sample were measured before each session (Thorlabs PM400). For Archon 1, t we used ∼24 W/mm² of 639 nm excitation. For Ace-mNeon, we used 100 mW/mm^2^ of 488 nm excitation. Fluorescence was collected on a scientific sCMOS camera (Hamamatsu ORCA-Fusion) at 500 frames/s.

### In vivo voltage imaging of corneal terminals and chemical stimulation

Mice were anesthetized (ketamine–domitor–saline 1:1:8, 12.5 µl/g, i.p.) and placed on a 37 °C heating pad; the head was fixed via the implanted head-plate. A custom eye-cup stabilized the eye (**Supplementary Fig. 1**) and was filled with SES that contained (in mM): 145 NaCl, 5 KCl, 2 CaCl₂, 1 MgCl₂, 10 glucose, 10 HEPES; pH 7.4; osmolarity ≥300 mOsm. MgCl₂ was added from a 1 M stock. Terminal arbors were identified by EGFP/mNeonGreen fluorescence. For examining the response to capsaicin or menthol, the most superficial (epithelium-proximal) terminal was selected; for spontaneous-activity recordings, the brightest terminal was selected. The protocol was 10 s of baseline recording, a 0.7-s puff, and 20 s of post-puff recording. Puffs were delivered from a pulled glass pipette (Sutter BF150-86-10; P-1000 puller) positioned 2–5 µm above the corneal surface and ∼10 µm laterally from the terminal, via a Picospritzer. Capsaicin (1 mM stock in ethanol) was diluted to 500 nM in SES; menthol (2 mM stock in ethanol) was diluted to 1 mM in SES; sulforhodamine-101 (SR101) was included as a visual marker.

### Image processing, signal extraction, and event detection

Raw movies were background-subtracted (constant offset) and photobleach-corrected (single-exponential fit to the whole-field mean intensity) prior to signal extraction. Videos were then high-pass-filtered in time with a 50-Hz high-pass filter, and segmented semi-automatically using principal component analysis followed by time-domain independent component analysis (PCA/ICA)^64^. The independent component corresponding to spiking activity was manually selected, and the spatial masks from PCA/ICA were then applied to the videos without high-pass filtering to extract fluorescence traces. The “before” trace was taken as the raw ICA time series, baseline corrected by subtracting a slow residual drift trend (sliding median, ∼3 s window) and expressed as an SNR-normalized signal: a binary mask excluded ±3 frames (∼±6 ms) around each detected spike; the noise standard deviation σ was computed over all subthreshold (unmasked) samples (N−1 estimator); and the normalized trace was signal/σ, so amplitudes are in multiples of baseline noise. Spikes were detected on a high-pass filtered version of the trace (100 Hz cutoff) by a threshold of 3× the filtered trace’s standard deviation, with the automatically detected threshold confirmed or manually adjusted by the experimenter.

Recordings after pharmacological blockade that contain no spikes and therefore cannot be demixed by ICA, whose spatial filter is estimated from spike-driven pixel covariance. For these, the terminal footprint was inherited from the paired before recording: its ICA spatial filter was binarized, registered onto the drug condition field of view using the best-bounded translation of the mean images, and the drug condition movie was projected through this mask as an equal-weight (unweighted) average. The “after” trace was baseline-corrected identically and normalized by the σ of the paired before movie, which was put through the same unweighted mask projection, not by its own σ, since block collapses the after-trace variance toward zero.

### Waveform, firing-rate and pattern analysis

Ongoing firing rate was computed per terminal as the total number of detected spikes divided by the duration of the analyzed recording and is reported in Hz; no minimum spike count was imposed beyond the single spike required for a terminal to be classified as active. Inter-spike-interval variability was quantified per terminal as the coefficient of variation of the inter-spike intervals (standard deviation divided by mean), computed only for terminals with at least two intervals. Per-spike waveforms were extracted in a 141-frame window (±70 frames, ≈±139.5 ms) centered on the peak (frame 71), baseline corrected (subtracting the mean of the first quarter of the window) and peak normalized per spike, then averaged per terminal and peak normalized again (time axis (−70:70)×1.993 ms). Half-width was obtained from a smoothing spline fit at the ≈0.5 level (threshold 0.51); afterhyperpolarization amplitude was the deepest post-peak value below baseline (0 if the waveform never dips below baseline); AHP area (AHP AUC) was the trapezoidal integral of the below-baseline post-peak segment (0 if no AHP). Waveform clustering used k-means (K = 3; 100 replicates; MaxIter 500; rng(123); squared Euclidean distance, the MATLAB default) on per-terminal peak-normalized waveforms. Firing patterns (Single/Burst/Mix) were classified in two stages: a one vs two component Gaussian mixture fit to the per-terminal ISI distribution (linear and log ISI space; acceptance of two components required ΔBIC, weight, count, separation, valley-to-peak, and mean ratio criteria), followed by a within-train dense/isolated spike analysis with a burst ISI threshold set at the GMM valley only when that valley exceeds the 70 ms a priori cutoff by less than 20 ms, else 70 ms. Terminals were labeled Single (one ISI cluster or fracSingle > 0.8), Burst (fracSingle < 0.2), or Mix (otherwise).

### Active and inactive terminals, and terminal brightness

A terminal was classified as “active” if the event-detection pipeline described above returned at least one spike in its ongoing-activity recording, and “inactive” if a terminal arbor was successfully imaged under identical conditions but no spike was detected in the recording.

Terminal brightness was measured as a per-frame fluorescence contrast: for each frame the mean intensity of pixels exceeding z = 1.645 (the upper 5% of the frame intensity distribution) minus the mean intensity of the lowest 5% of pixels, averaged over the first 2,500 frames (5 s) of the recording, with spike-containing frames excluded for active terminals and blank or saturated frames excluded for inactive terminals.

Because a terminal too dim to yield a detectable signal cannot be scored as inactive, active and inactive sets were brightness-matched before prevalence was computed: inactive terminals whose brightness fell below the dimmest active terminal (151.1 camera counts for Archon1) were excluded, leaving 110 active and 142 inactive Archon1 terminals, so that the reported prevalence (43.7%) refers only to the brightness range in which ongoing activity was ever observed. The same procedure was applied to the Ace-mNeon2 dataset (threshold 126.3 camera counts), Within active terminals, the association between brightness and ongoing firing rate was tested by Pearson correlation (inactive terminals were assigned a firing rate of zero by definition and were therefore excluded from the correlation and are shown in the scatter for reference only).

### Pharmacological experiments

Oxybuprocaine (0.4%, 0.5 µl) was applied directly to the eye; after 3 min, ∼1 ml SES was added to the eye-cup and recordings resumed. The drug was then washed out with SES and wash-out effects assessed ∼30 min later^17^.

Suzetrigine (VX 548, MedChemExpress) was dissolved to a 25 mM stock in DMSO (2.36 mg/200 µL) and diluted directly in SES to 0.1 mM working solutions applied to the eye cup for 15 min before recording^46^.

XE991 dihydrochloride (Kv7/KCNQ blocker, R&D Systems, MW 449.36 g/mol) was dissolved to a 7.52 mM stock in DDW (3 mg / 0.888 mL) and diluted fresh in SES to a working concentration (50 µM or 0.376 mM) on the day of recording, applied to the eye cup for 15 min before recording^17^.

ZD7288 hydrate (Sigma Z3777, HCN/Ih blocker, MW 292.81 g/mol) was dissolved to a 10 mM aqueous stock in DDW and diluted fresh in SES to the working concentration of 5 mM on the day of the experiment, applied to the eye cup for 15 min before recording^48^.

### Lidocaine delivery to the trigeminal ganglion

The cannula insertion for lidocaine delivery to the TG was performed as previously explained^65^. In brief, under ketamine–domitor–saline anesthesia, the scalp was cleaned (Betadine), the head stabilized, and a single cannula (RWD Brain Single Cannula) lowered to −6.2 mm through the pre-drilled injection hole and fixed with dental cement; a dummy cap was inserted until use. For blockade, 2% lidocaine mixed with Evans Blue (delivery marker) was injected (up to 4 µl; 20 nl/s, UMP3 microinjection pump, World Precision Instruments)). Voltage imaging was performed before and 3 min after delivery. After imaging, the TG was extracted and inspected for blue coloration to confirm delivery.

### Behavioral capsaicin-sensitivity assay

Assessment of sensitivity to capsaicin instillation was performed as previously described^17^. In short, capsaicin (500 nM in 1 ml) was instilled onto the cornea and forelimb eye-wipes counted per 5 min before and after instillation, comparing AAV9-EGFP and AAV9-Archon1-EGFP mice.

### Statistics

Analyses were performed in MATLAB (MathWorks) and Prism 11 (GraphPad). Quantitative data were expressed in text as the Mean ± SD. Boxplots presented in the Figures depict the Median, 25^th^ and 75^th^ percentiles, interquartile range (IQR), and min and max values within 1.5*IQR (whiskers). Sample sizes were based on prior studies using related *in vivo* corneal imaging approaches^17^. Mice were randomly allocated to groups in all experiments. Normality was assessed using the Shapiro-Wilk test. For normally distributed values, two-tailed unpaired Student *t*-tests, paired *t*-tests, ordinary one-way analysis of variance (ANOVA), two-way/mixed ANOVA, and Repeated-measures (RM) ANOVA were used when appropriate. For values that were not normally distributed (Fig. 4-6, Supplementary Fig. 4), Wilcoxon signed-rank and Kruskal-Wallis tests were used as appropriate. Actual *p* values are presented for each data set. The criterion for statistical significance was p < 0.05.

The independent unit of analysis was a single terminal; when two terminals belonged to the same axon, one was selected at random and the other excluded (except in Fig. 4, where sister terminal trees and the parent axon were deliberately recorded together for the terminal–axon comparison). For the paired and grouped comparison Figures (apart from Figure 2, Supplementary Figs. 5 and 6) in which data from several terminals from the same eye were collected, each plotted data point corresponds to one terminal recorded from one eye of one mouse, so that terminal, eye, and animal are equivalent units for those comparisons. Exact tests, n (terminals and mice), and P values are given in each Figure and figure legend.

## Data and code availability

All data needed to evaluate the conclusions in the paper are present in the paper and/or the Supplementary Materials. Further information and requests for source data for each panel in main and supplementary figures, resources, and reagents should be directed to and will be fulfilled by the Lead Contacts, Alexander Binshtok and Yoav Adam

## Acknowledgments

We thank Maya Groysman and the ELSC vector core facility (EVCF) team for virus production, and Itamar Frachtenberg and the ELSC Fab Lab team for help with the design and fabrication of the custom eye caps.

## Author contributions

Conceptualization, A.B. and Y.A.; Investigation, E.S., R.B., S.L., S.B.S.; Formal Analysis, E.S., R.B., Y.A. and A.B.; Writing – Original Draft, Y.A. and A.B.; Funding Acquisition, A.B. and Y.A.; Supervision, A.B. and Y.A.

## Supplementary Figures and Figure Legends

**Supplementary Fig. 1.**
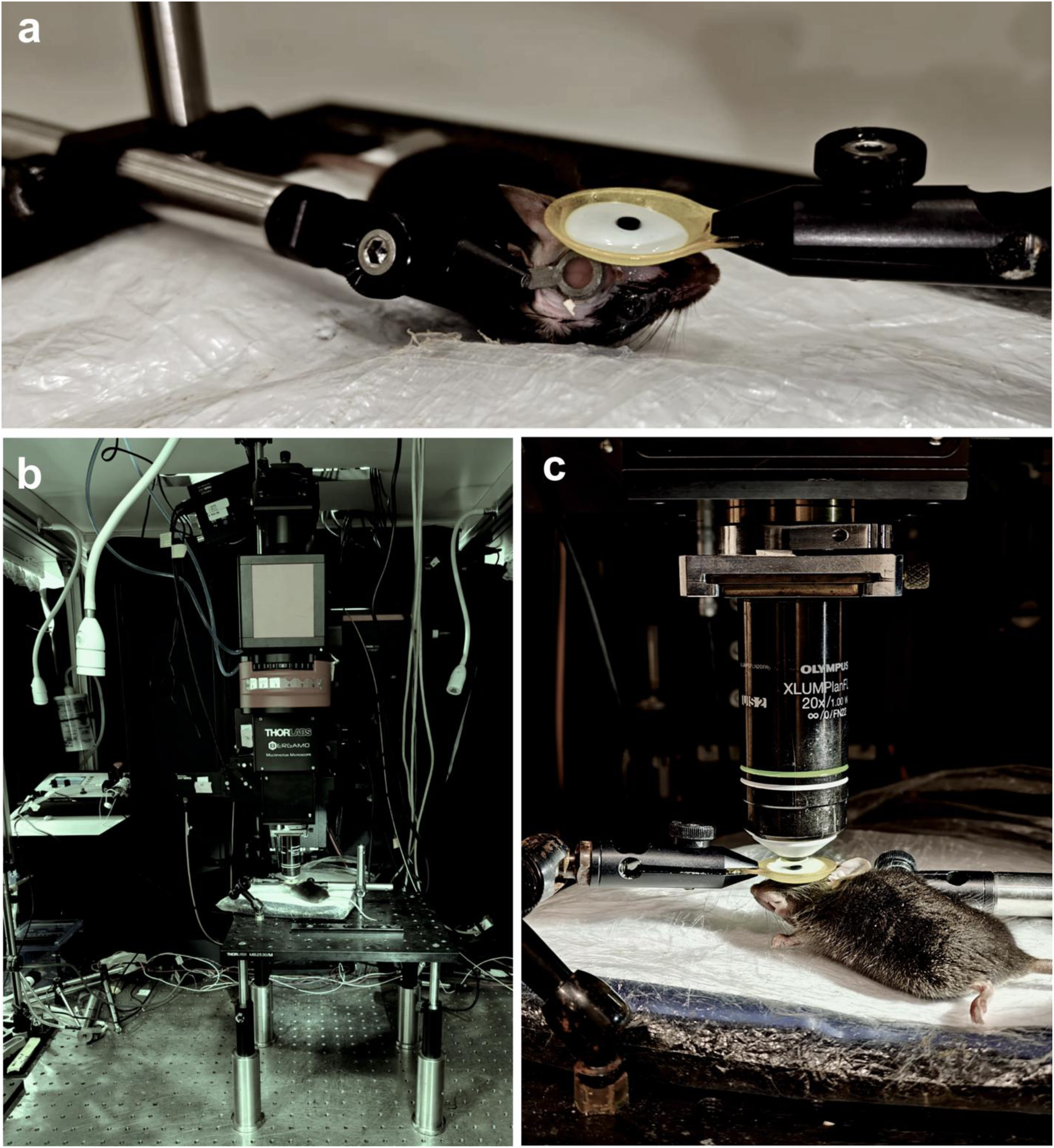
In vivo voltage-imaging preparation for recording from corneal nociceptive terminals. (a) Lateral close-up of an anesthetized mouse with the head stabilized and the eye engaged by a custom eye-cup (a silicone ring with a central aperture) held by an articulated arm; flanking arms provide additional fixation. The eye-cup is filled with standard extracellular solution, which keeps the cornea moist, permits incubation with pharmacological agents in a subset of experiments, and limits eye movement during imaging. (b) Overview of the imaging rig: an upright Thorlabs Bergamo widefield/multiphoton microscope mounted on a vibration-isolated optical table, with the animal platform positioned beneath the objective. **(c)** Front view of the animal under the objective (Olympus XLUMPlanFL N 20×/1.00 NA water-immersion), showing the objective aligned over the eye-cup and the eye; the mouse rests on a temperature-controlled heating pad on the translation stage, with the eye-cup holder and fixation arms in place. Images are representative of the preparation used throughout.

**Supplementary Fig. 2.**
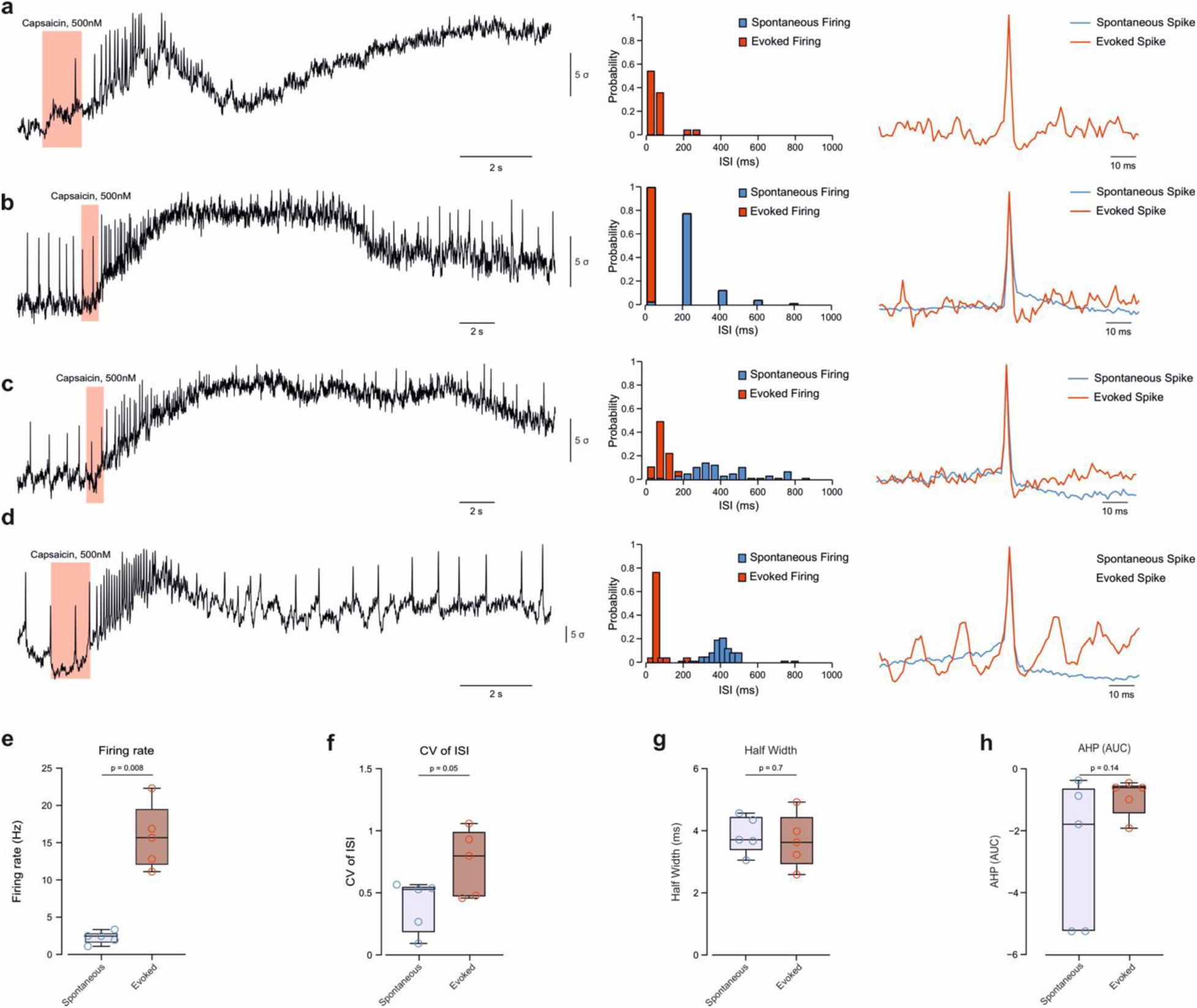
Spontaneous and capsaicin-evoked events share waveform properties but differ in rate (related to Fig. 1c). (**a–d**) Four representative terminals, each showing (*left*) a voltage trace with capsaicin application (500 nM, pink), (*middle*) ISI probability histograms for spontaneous (*blue*) and evoked (red) firing, and (*right*) overlaid mean spontaneous and evoked waveforms (scale 10 ms). Panel **a** illustrates a capsaicin-responsive terminal with little or no detectable spontaneous firing. (**e–h**) Paired spontaneous vs evoked comparison: (**e**) firing rate; (**f**) CV of ISI; (**g**) half-width; (**h**) AHP area. Box plots with individual terminals; paired t-test; n = 5 terminals from N = 5 mice.

**Supplementary Fig. 3.**
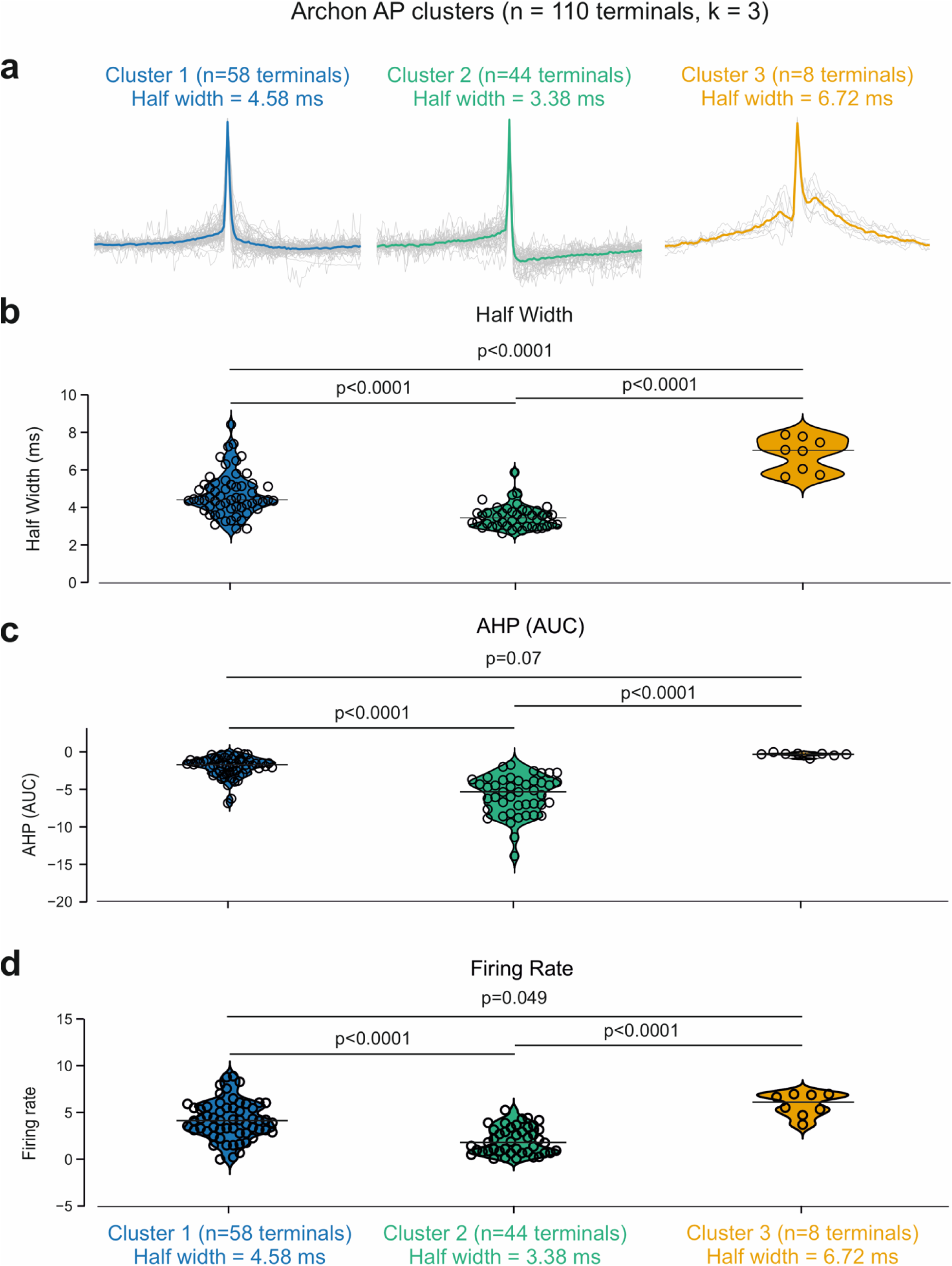
Three waveform clusters differ in shape and firing rate (related to Fig. 2). (a) Mean spike waveforms for three k-means clusters of Archon1 active terminals (n = 110 terminals; N = 42 mice, k = 3): cluster 1 (n = 58 terminals, half-width 4.58 ms), cluster 2 (n = 44 terminals, 3.38 ms), cluster 3 (n = 8, 6.72 ms). (b–e) Per-cluster distributions of half-width (b), afterhyperpolarization (c), AHP area (d) and firing rate (e), with pairwise two-sided Wilcoxon rank-sum P values (uncorrected). Note that half-widths were used to define the clusters; the firing-rate difference (e; cluster 2 vs 3 P = 0.049, others P < 0.0001) is the comparison independent of the clustering features.

**Supplementary Fig. 4.**
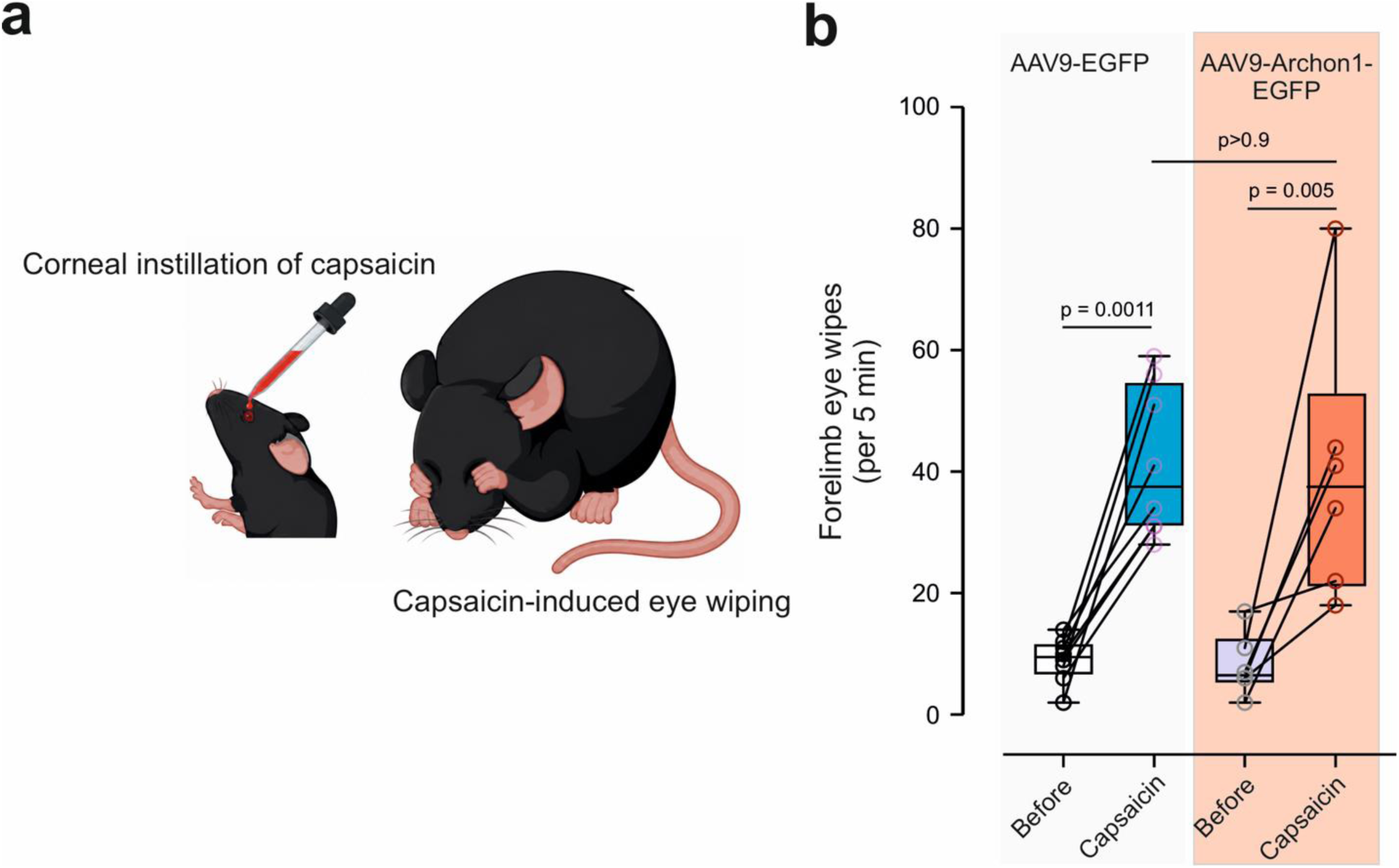
Archon1 expression does not detectably alter capsaicin-evoked nocifensive behavior (related to Fig. 2). (a) Schematic of the assay: capsaicin is instilled onto the cornea, and the resulting nocifensive response is quantified as forelimb eye-wiping. (b) Forelimb eye-wipes per 5 min before and after corneal capsaicin instillation, in mice expressing AAV9-EGFP (control; *left*) or AAV9-Archon1-EGFP (*right*). Capsaicin increased eye-wiping in both groups, and the capsaicin-evoked response did not differ between groups. Box plots show the median and interquartile range with whiskers; each circle is one mouse, and connecting lines indicate the same mouse before and after capsaicin. n = 8 mice (AAV9-EGFP) and 6 mice (AAV9-Archon1-EGFP), two-way mixed-design ANOVA (repeated measures on the before/after factor; the data are normally distributed), with Bonferroni post-hoc.

**Supplementary Fig. 5.**
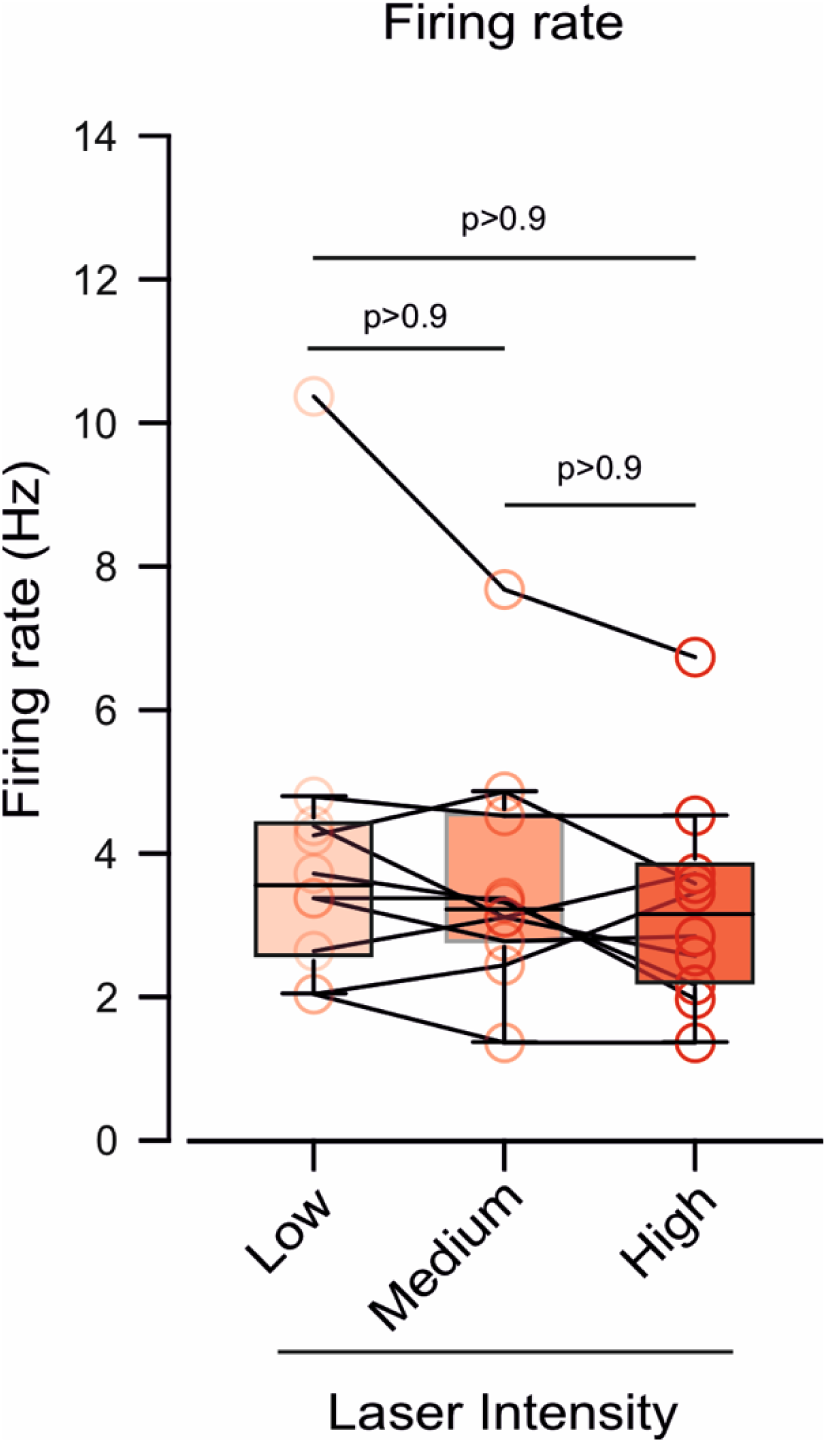
Ongoing firing rate is independent of excitation-laser power (related to Fig. 2). Firing rate at low, medium and high excitation-laser intensity for the same terminals (lines connect the same terminal); all pairwise, n = 10 terminals from N = 10 mice, Kruskal-Wallis test.

**Supplementary Fig. 6.**
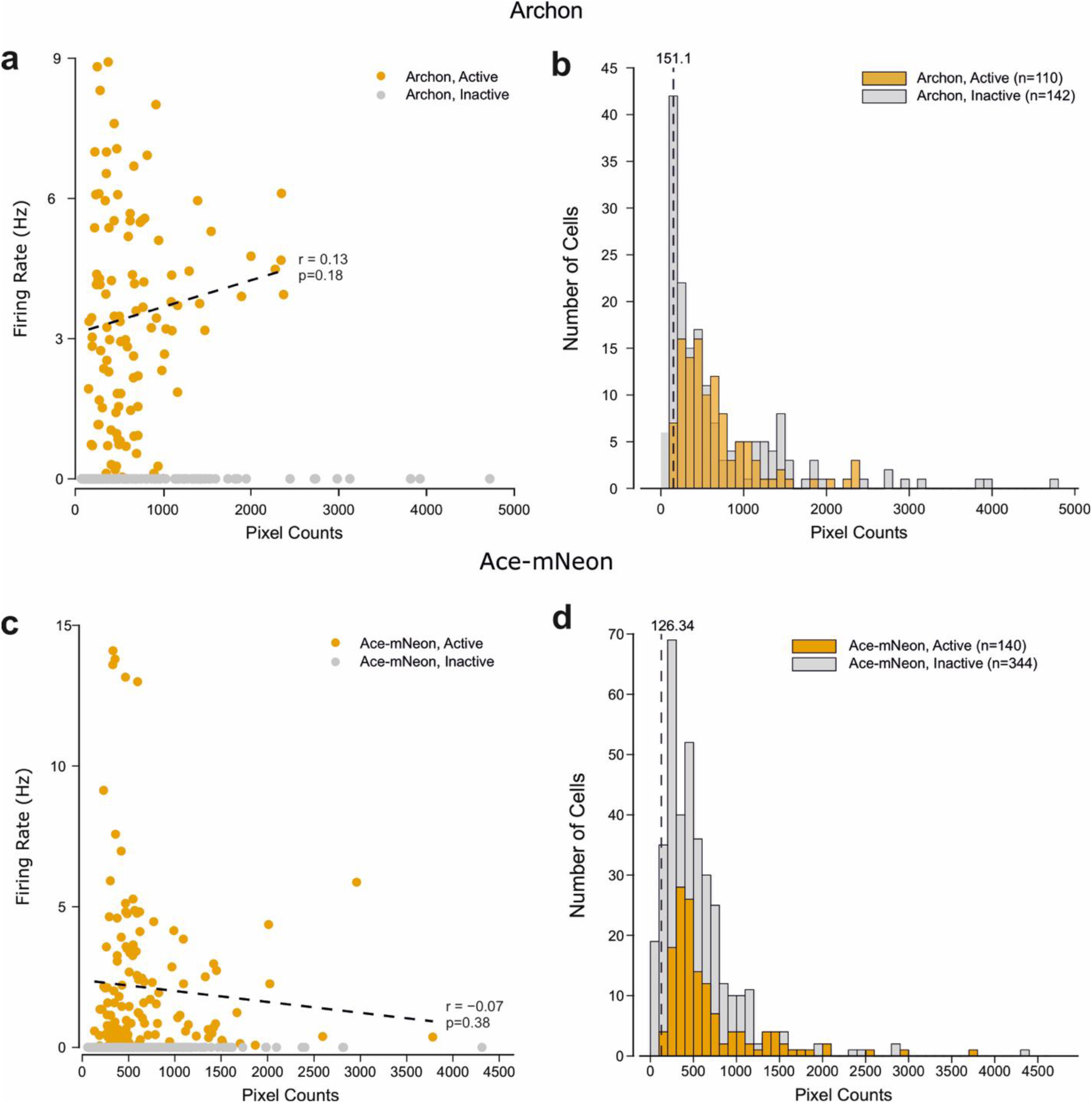
Ongoing firing is reproduced with a second indicator and is not explained by terminal brightness (related to Fig. 2). (a) Archon1: ongoing firing rate versus terminal brightness (integrated pixel counts) for individual terminals classified as active (terminals that demonstrate ongoing activity, *see Methods*, *orange*) or inactive (*grey*); firing rate was uncorrelated with brightness (r = 0.13, p = 0.18; dashed line, linear fit). (b) Distribution of terminal brightness for active (n = 110) and inactive (n = 142) Archon1 terminals; dashed line, minimal brightness among active terminals (151.1). (c) Ace-mNeon2: ongoing firing rate versus terminal brightness, classified as active or inactive; firing rate was uncorrelated with brightness (r = −0.07, P = 0.38). (d) Distribution of terminal brightness for active (n = 140) and inactive (n = 344) Ace-mNeon2 terminals; dashed line, minimal brightness among active terminals (126.34). Active and inactive terminals overlapped in brightness for both indicators, so ongoing firing was not distinguished by signal intensity. Each point/count is one terminal.

**Supplementary Fig. 7.**
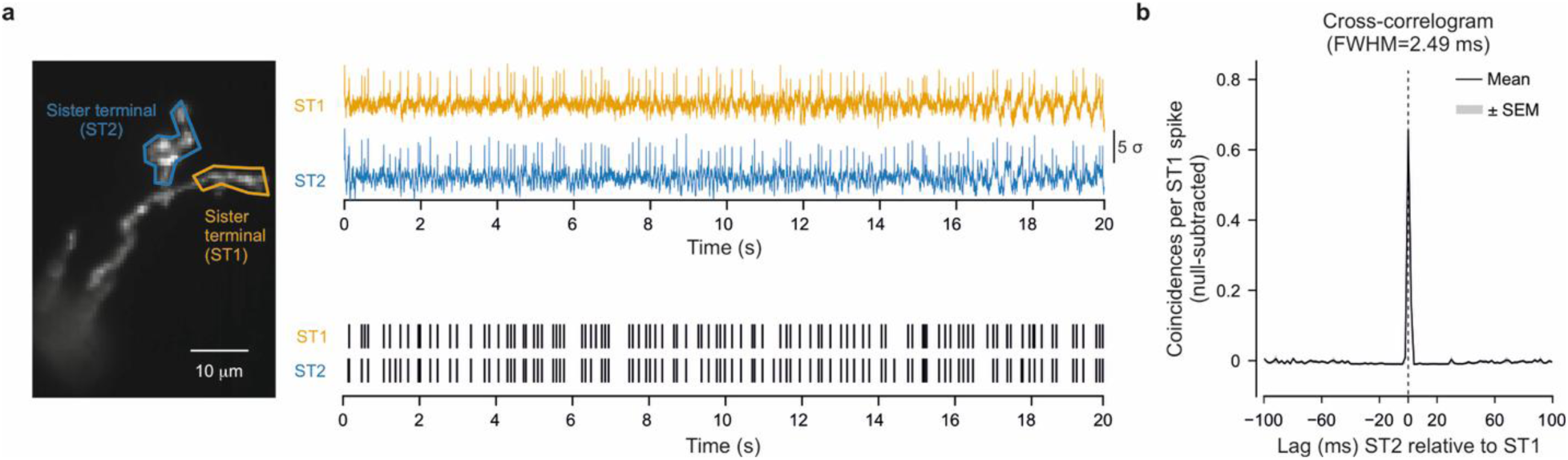
Sister terminals recorded with Ace-mNeon2 fire in tight synchrony (related to Fig. 4c). **(a)** *Left*: fluorescence image with two manually defined regions of interest, sister terminal 1 (ST1, *orange*) and sister terminal 2 (ST2, *blue*. *Right*, top: simultaneously recorded voltage traces from ST1 (*orange*) and ST2 (*blue*); bottom: corresponding spike rasters for ST1 and ST2, showing near-coincident firing. **(b)** Pairwise cross-correlogram (mean ± SEM across pairs) of null-subtracted coincidences per ST1 spike as a function of lag (ST2 relative to ST1). The correlogram peaks sharply at zero lag (FWHM of 2.49 ms) with no consistent lead or lag, indicating near-simultaneous firing with no detectable directional offset. Data are from n = 5 sister-terminal pairs from N = 5 mice.

---

**Supplementary Movie 1.** Three-dimensional reconstruction of a corneal nociceptive terminal arbor and its parent axon (related to Fig. 4a,b) Rotating volume rendering of an *in vivo* two-photon *z*-stack of a single Ace-mNeon2-labeled peripheral terminal arbor innervating the cornea. The parent axon can be followed as it ascends through the tissue and branches into the terminal arbor and its individual terminal branches, illustrating its three-dimensional organization. Acquisition: z-stack of 107 optical sections at 0.5 µm steps spanning 53.5 µm in depth.

